# How biological agents can couple neural task modules for dealing with the stability-plasticity dilemma

**DOI:** 10.1101/457150

**Authors:** Pieter Verbeke, Tom Verguts

**Affiliations:** Department of experimental psychology; Ghent University; B9000 Belgium

## Abstract

We provide a novel computational framework on how biological and artificial agents can learn to flexibly couple and decouple neural task modules for cognitive processing. In this way, they can address the stability-plasticity dilemma. For this purpose, we combine two prominent computational neuroscience principles, namely Binding by Synchrony and Reinforcement Learning. The model learns to synchronize task-relevant modules, while also learning to desynchronize currently task-irrelevant modules. As a result, old (but currently task-irrelevant) information is protected from overwriting (stability) while new information can be learned quickly in currently task-relevant modules (plasticity). We combine learning to synchronize with several classical learning algorithms (backpropagation, Boltzmann machines, Rescorla-Wagner). For each case, we demonstrate that our combined model has significant computational advantages over the original network in both stability and plasticity. Importantly, the resulting models’ processing dynamics are also consistent with empirical data and provide empirically testable hypotheses for future MEG/EEG studies.

**Author summary:** Artificial and biological agents alike face a critical trade-off between being sufficiently adaptive to acquiring novel information (plasticity) and retaining older information (stability); this is known as the stability-plasticity dilemma. Previous work on this dilemma has focused either on computationally efficient solutions for artificial agents or on biologically plausible frameworks for biological agents. What is lacking is a solution that combines computational efficiency with biological plausibility. Therefore, the current work proposes a computational framework on the stability-plasticity dilemma that provides empirically testable hypotheses on both neural and behavioral levels. In this framework, neural task modules can be flexibly coupled and decoupled depending on the task at hand. Testing this framework will allow us to gain more insight in how biological agents deal with the stability-plasticity dilemma.

## Introduction

Humans and other primates are remarkably flexible in adapting to constantly changing environments. Additionally, they excel at integrating information in the long run to detect regularities in the environment and generalize across contexts. In contrast, artificial neural networks (ANN), despite being used as models of the primate brain, experience significant problems in these respects. In ANNs, extracting regularities requires slow, distributed learning, which does not allow strong flexibility. Moreover, fast sequential learning of different tasks typically leads to (catastrophic) forgetting of previous information (for an overview see (1)). Thus, ANNs are typically unable to find a trade-off between being sufficiently adaptive to novel information (plasticity) and retaining older information (stability), a problem known as the stability-plasticity dilemma. Additionally, it remains unknown how biological agents deal with this dilemma.

We provide a novel framework on how biological and artificial agents may address this dilemma. We combine two prominent principles of computational neuroscience, namely Binding by Synchrony (2–5) and Reinforcement Learning (RL; 6,7). In BBS, neurons are flexibly bound together by synchronizing them via oscillations. This implements selective gating (e.g., 8) in which synchronized neurons communicate efficiently, while desynchronized neurons do not. Thus, BBS allows to flexibly alter communication efficiency on a fast time scale. By using RL principles, the model can learn autonomously which neurons need to be (de)synchronized.

In the modeling framework, BBS binds relevant neural groups, called (neural task) modules, and unbinds irrelevant modules. This causes both efficient processing and learning in synchronized modules; and inefficient processing and learning in desynchronized modules. The resulting model deals with the stability-plasticity dilemma by flexibly switching between task-relevant modules and by retaining information in task-irrelevant modules. An RL unit (9) uses reward prediction errors to evaluate whether the model is synchronizing the correct task modules. We apply our framework to networks that themselves learn via three classic synaptic learning algorithms, namely backpropagation (10), Restricted Boltzmann machines (RBM; 11) and Rescorla-Wagner (RW; 12,13).

The model consists of three units (Figure 1A). The Processing unit contains a network consisting of a number of task-specific modules. In addition, RL and Control units together form an hierarchically higher Actor-Critic structure, modeled after basal ganglia/primate prefrontal cortex (14). The RL unit (modeling ventral striatum/ anterior medial frontal cortex) evaluates behavior. More specifically, it learns to assign a value to a specific task module (how much reward it receives by using this module) and compares this value with the externally received reward to compute prediction errors. Additionally, the RL unit has a Switch neuron (see Figure 1C and D). This Switch neuron computes a weighted sum of negative prediction errors over trials. When this sum reaches a threshold of .5, it signals the need for a strategy switch to the Control unit (see Methods for details). This Control unit functions as an Actor in order to drive neural synchronization in the Processing unit. One part of the Control unit (modeling lateral frontal cortex (LFC)) contains task units that point to task modules in the Processing unit (15); another part (modeling posterior medial frontal cortex (pMFC)) synchronizes task modules based on those task units (16). Crucially, LFC and pMFC both use prediction error information, but on different time scales. While the LFC uses prediction errors on a slow time scale to know when the task rule has changed and a switch of modules is needed, the pMFC uses prediction errors on a fast time scale to enhance control over the synchronization process as soon as a negative prediction error occurs.

**Figure 1.**
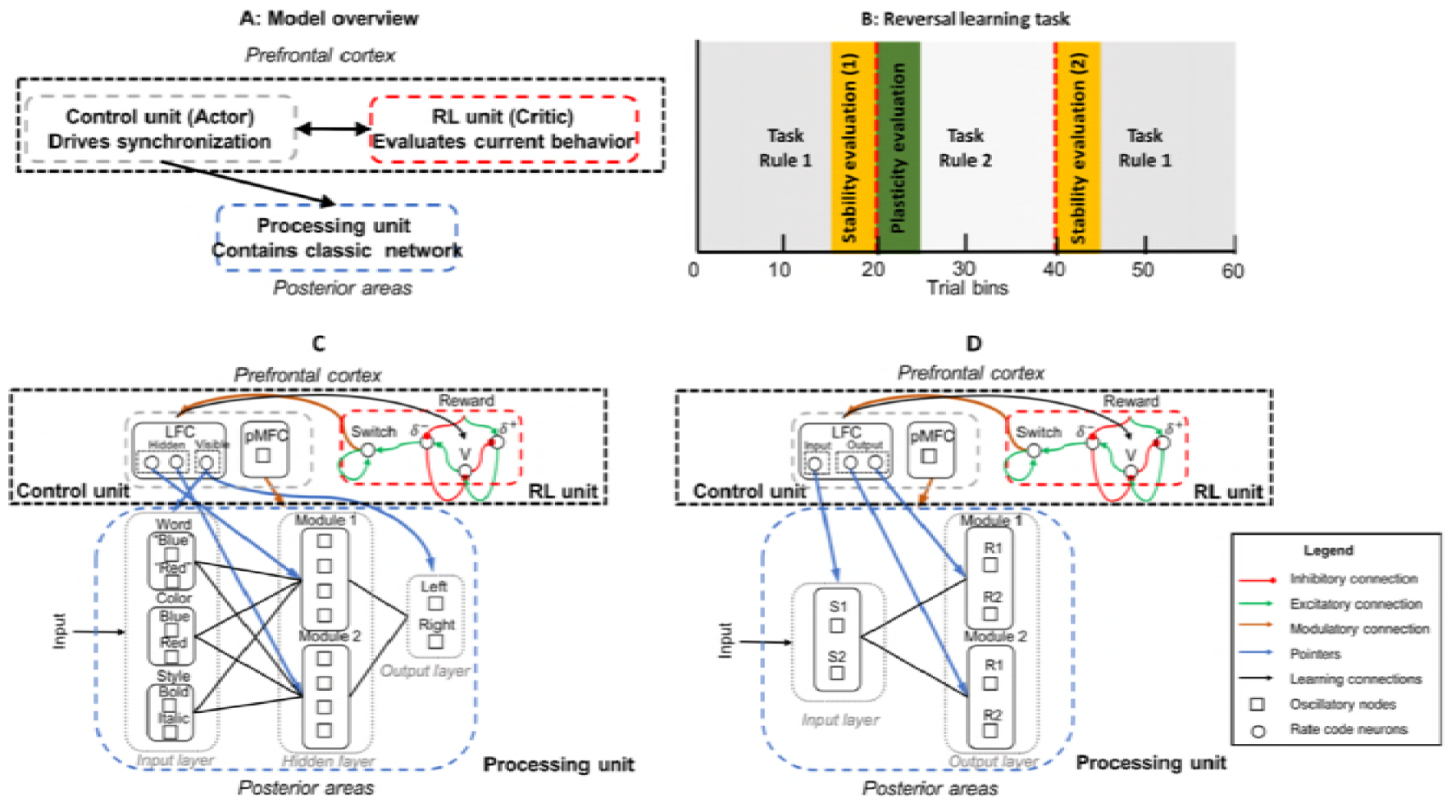
Model and task overview. **A:** General model overview. **B:** Reversal learning task. Trial bin size = 40 trials for multi-layer models and trial bin size = 4 trials for models with RW networks (see Methods for details). Red dotted vertical lines indicate task switches. **C:** More detailed overview of the multi-layer model in the context of a Stroop task. **D**: More detailed overview of the RW model in the context of an S-R associative learning task.

In order to drive neural synchronization we rely on the idea of binding by random bursts (16–18). Here, applying positively correlated noise to two oscillating signals reduces their phase difference. In addition to implementing binding by random bursts, the current work also implements unbinding by random bursts. In particular, applying negatively correlated bursts increases the phase difference between oscillating signals and thus unbinds (i.e., dephases) the two signals.

We test our model on a (cognitive control) reversal learning task. Here, each hierarchically lower algorithm (e.g., Boltzmann) sequentially learns different task rules. The relevant task rule changes during the task (Figure 1B). The model must detect when task rules have changed, and flexibly switch between different rules without forgetting what has been learned before. Our task is divided in three equally long blocks that alternate between two task rules (rule 1-rule 2-rule 1). For the backpropagation and RBM networks (because of their hidden layer further called multi-layer networks), a multiple-feature Stroop-like task is used. Here, stimuli are presented that contain three crucial features. They are words (“red” or “blue”) printed in a certain color (red or blue) and style (bold or italic). There are two response options. The task is to respond to the word when it is printed in bold and to the color when it is printed in italic. During rule 1 they should respond with Response 1 (R1) for red and Response 2 (R2) for blue. This is reversed for rule 2. For the RW network, which cannot handle such complex task rules, we use simple Stimulus-Response (S-R, linearly separable) associations. According to rule 1, R1 leads to reward after presentation of Stimulus 1 (S1) and R2 leads to reward after presentation of Stimulus 2 (S2). For rule 2 these associations are reversed, linking R1 with S2 and R2 with S1. The Stroop-like task consisted of 2400 trials and the S-R associative learning task of 240 trials. For comparison, we divided them in 60 trial bins for some analyses and plots. Figure 1C and D illustrate the detailed model build-up in respectively the Stroop-like task and the S-R associative learning task. We compare our combined (henceforth, full) models with models that only use synaptic learning (i.e., only contain the Processing unit; called synaptic models). We evaluate plasticity as the ability to learn a new task after learning a different task; and stability as the interference of learning a new task on performance on the old task (see Figure 1B and Methods).

## Results

### The stability-plasticity dilemma

#### Backpropagation

Figure 2A-C show a clear advantage for the full relative to the synaptic backpropagation model in overall accuracy as well as plasticity and stability. This advantage was present across all learning rates. This advantage appears because the synchronization supports modularity, thus protecting information from being overwritten.

**Figure 2.**
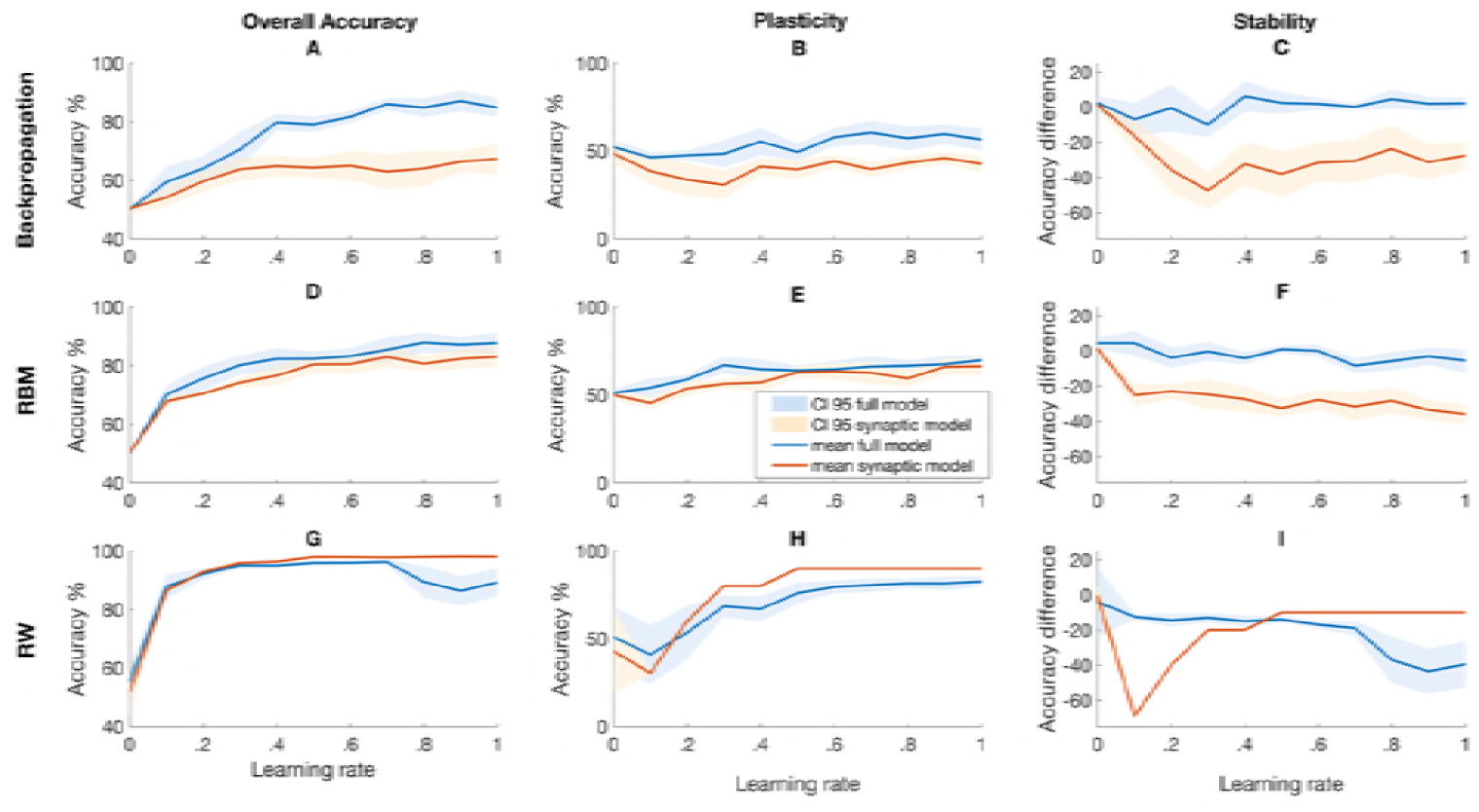
Performance of models on reversal learning task. Blue lines show means for the full model and orange lines represent the mean values for the synaptic models. The shades indicate the corresponding 95% confidence intervals.

#### RBM

Also panels D-F of Figure 2 show an advantage for the full model relative to the synaptic RBM model. This advantage is less strong than for the backpropagation model because the synaptic RBM model shows a stronger plasticity than the synaptic backpropagation model.

#### RW

Figure 2G-I shows similar overall accuracy for the full and synaptic RW models. When synaptic learning rates are slow (β = .1-.4), the full model has a better stability than the synaptic model. However, this advantage disappears for higher learning rates and the synaptic model shows a higher plasticity than the full RW model. There is also a dip in performance for higher learning rates in the full RW based model. Reasons for this dip are explained in the Methods section.

### Model dynamics

More insight into the dynamics of the model is given in Figure 3. We show data for simulations with a learning rate of .3.

**Figure 3.**
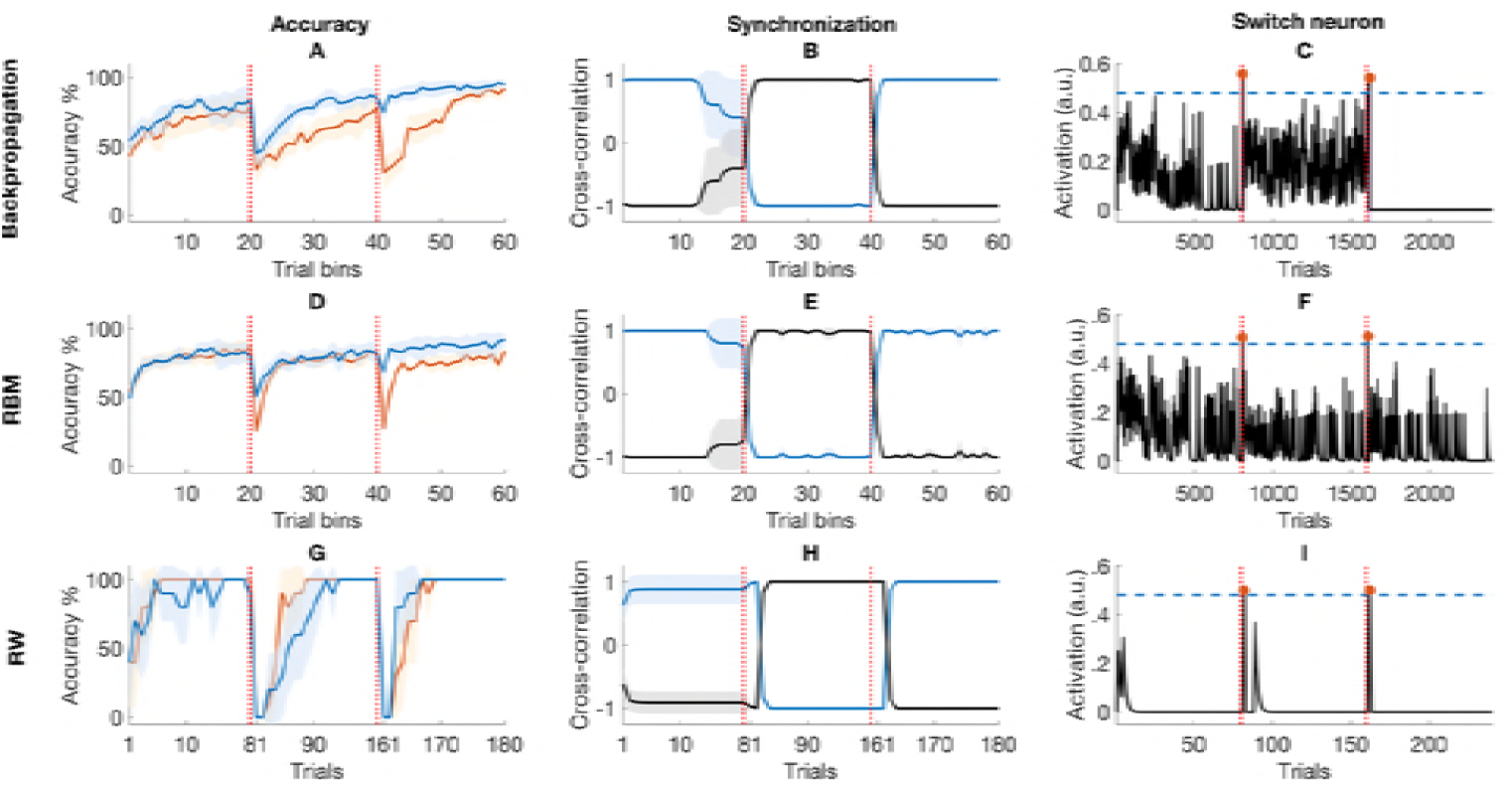
Model data. Model dynamics are shown for simulations with a learning rate of .3. In column 1 (panels A, D and G), the blue lines represent data of the full model and the orange lines represent data for the synaptic model. In column 2 (panels B, E and H), blue lines represent values for the initially (randomly) chosen module and black lines for the other module. In column 3 (panels C, F and I), activity of the Switch neuron (see Figure 1C and D) is shown for one selected simulation of the model (in black). Blue horizontal dashed lines indicate the threshold of the Switch neuron and the orange dots indicate data points above the threshold. In all panels, red vertical dotted lines indicate task switches and shades indicate 95% confidence intervals.

#### A closer look at accuracy

Figure 3A illustrates the accuracy evolution over the whole task for both the full and synaptic backpropagation model. During the first part of the task, the synaptic and full model show a similar performance. When there is a first switch in task rule, the drop in accuracy is slightly larger for the synaptic model than for the full model. This is caused by the fact that the synaptic model has to learn task rule 2 with weights that were pushed in the opposite direction during learning of task rule 1. Instead, the full model switches to another task module and starts learning from a random weight space. After the second rule switch, there is again a strong decrease of accuracy in the classic model but not in the full model. Here, the classic model had to relearn the first task rule (catastrophic forgetting) while the full model switched to the first module where all old information was retained. As illustrated in Figure 3D, these findings are replicated by the RBM model. In Figure 3G, the accuracy is plotted for the RW model. As suggested by Figure 2G-I, the full model shows a similar performance during the first part of the task, a lower plasticity after the first task switch but a higher stability after the second task switch compared to the synaptic model.

#### Synchronization of modules

Figure 3B represents the synchronization between the input layer and different task modules for the backpropagation model. Here, we see that the model performs quite well in synchronizing task-relevant and desynchronizing task-irrelevant modules. Additionally, the model is able to flexibly switch between modules. A similar pattern is observed in Figure 3E and Figure 3H where the data for respectively the RBM and RW models are shown. In these plots, we observe wider confidence intervals in some trial bins. This reflects the fact that the model sometimes also erroneously switches. However, if such an incorrect switch occurs, the model will also switch back to the correct module.

#### The Switch neuron

Figure 3C show activation in the Switch neuron for the backpropagation model. Crucially, we observe in this plot only two points above the threshold of .5. These two points are right after the first task rule switch and right after the second task rule switch. Thus, the model correctly decides when a switch is necessary. A similar phenomenon occurs in the RBM model (Figure 3F) and the RW model (Figure 3I). The exact dynamics of the Switch neuron are most clearly observed for the backpropagation model (Figure 3C). During the first trials of learning a new task rule (approximately trials 1-200 and 800-1000) a new task module is used. Typically, a new module has not learned anything and makes a lot of errors. Here, the high number of (prediction) errors during learning is reflected in a constant high activation of the Switch neuron during these trials. However, a newly used task module also starts with a low predicted value (variable *V*, see equation (7) in Methods) and hence every error only elicits a small negative prediction error which is not enough for the Switch neuron to reach the threshold. When the task module learns the task, it produces less errors but it also learns to assign a high value to that module, resulting in stronger prediction errors when an error occurs. Hence, in later trials there is a weaker mean activation in the Switch neuron of almost zero, with occasional strong bursts of activity when a rare prediction error does occur. However, also this is typically not enough to reach the Switch threshold. The Switch threshold is only reached when a module that has a high value assigned to it, makes several consecutive errors. This typically means that the module is used in the wrong context and hence a switch is needed. After the second task switch, activation in the Switch neuron of the backpropagation model (Figure 3C) remains at zero. This is because the model reached full convergence and makes no errors anymore. This is not observed in the RBM model (Figure 3F) because it uses a probabilistic response threshold, making the model always susceptible to a small number of errors. Finally, activation of the Switch neuron for the RW model (Figure 3I) after every switch almost immediate converges towards zero, indicating that the RW learning algorithm is very fast and efficient.

### Connections with empirical data

As a model of how the brain controls its own processing, we next aimed at connecting with empirical data and describe testable hypotheses for future empirical work. For reasons described in the Methods section we only present data for the backpropagation and RW model here.

#### Feedback Related Negativity

As described in the Methods section, theta amplitude in the pMFC gradually decayed during the whole task. However, when a negative prediction error occurred the pMFC network node received a burst which increased its amplitude again. This can be clearly observed in the ERP that is plotted in Figure 4A for the backpropagation model and Figure 4D for the RW model. Here, the bursts occurring from approximately 100 to 300 ms after feedback results in a strong negative peak around 200 msec, corresponding to the empirical feedback related negativity (FRN; e.g., 19–24).

**Figure 4.**
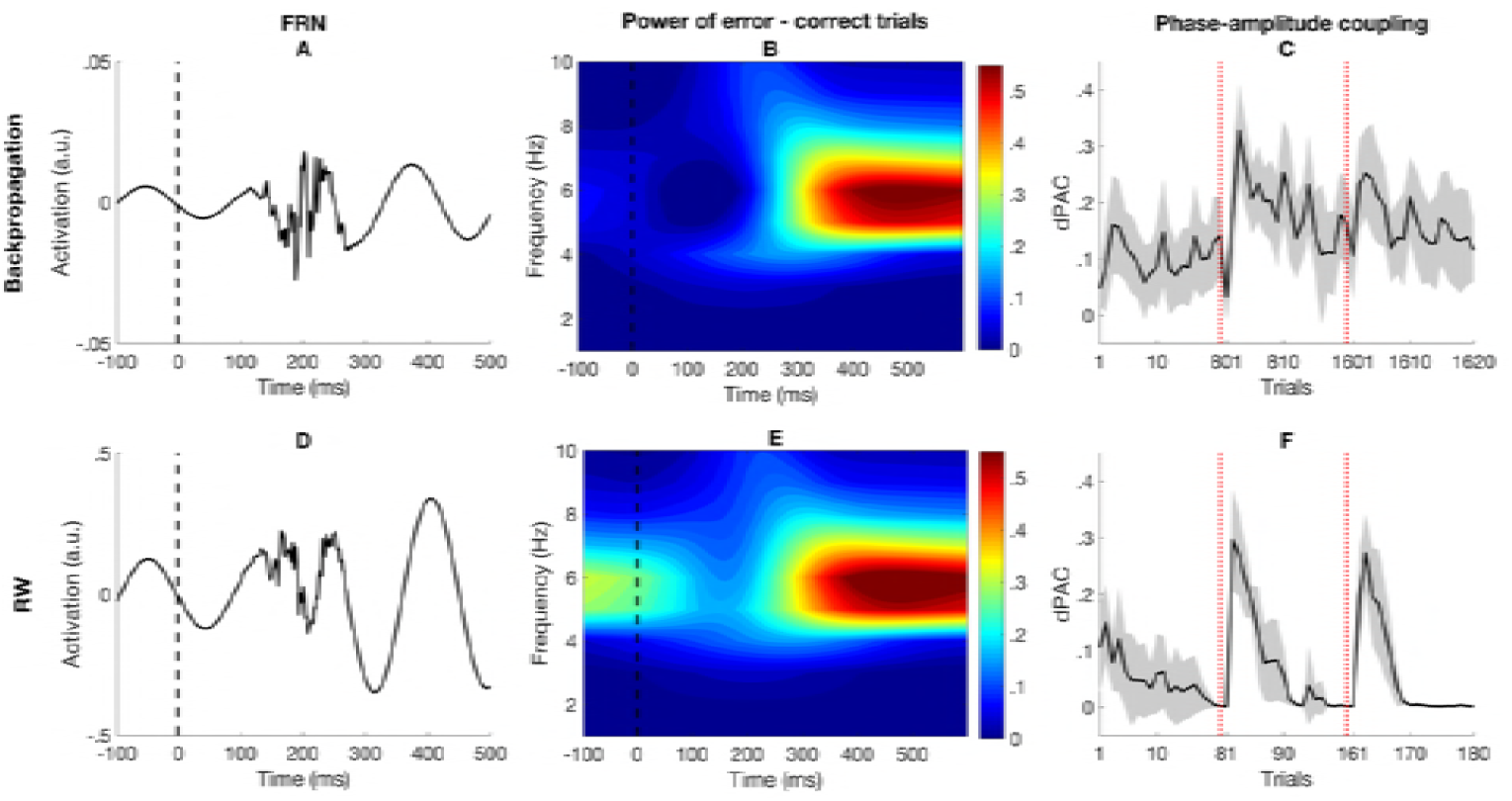
Model predictions for empirical data. Black vertical dashed lines indicate the moment of reward feedback. Red vertical dotted lines indicate task switches. Shades illustrate 95% confidence intervals.

#### Theta power

Additionally, we performed time-frequency decomposition of the signal produced by the pMFC node. More specifically, we were interested in theta power after feedback. We computed the contrast of power in the inter-trial interval after error and after correct trials in the time-frequency domain. Also here, and in accordance with previous empirical work (e.g., 24–26), we clearly observe increasing theta power, starting 200 ms after negative feedback. Again, this is shown both for the backpropagation (Figure 4B) and RW (Figure 4E) model.

#### Phase-amplitude coupling

Figure 4C, F illustrate the coupling between the phase of theta-oscillations in the pMFC and gamma amplitude in the Processing unit. Again consistent with empirical data (27,28), these plots show a clear increase in phase-amplitude coupling after a task rule switch. This is mainly caused by the fact that there are many negative prediction errors in these trials. These prediction errors increase theta power in the pMFC which in turn increases the number of bursts received by the gamma oscillations in the Processing unit (see Methods). This combination of events results in an increase of phase-amplitude coupling (PAC).

## Discussion

We described a computationally efficient and biologically plausible framework on how biological and artificial agents may deal with the stability-plasticity dilemma. We combined two neurocomputational frameworks, BBS (2–4) and RL (6). BBS flexibly (un)binds (ir)relevant neural modules and RL autonomously discovers which modules need to be (un)bound. Thus, the model could flexibly switch between different tasks (plasticity) without catastrophically forgetting older information (stability). We demonstrated that the model was consistent with several behavioral and electrophysiological (e.g., EEG/MEG) data.

Our model consists of three units. The Processing unit contains a task-processing network, trained by a classical learning rule (backpropagation, RBM or RW). Anatomically, it can be localized in several posterior (neo-)cortical processing areas, depending on the task at hand. Its activity is strongly stimulus-dependent and synaptic strengths change slowly. The RL unit learns to attach value to specific task modules, based on prediction errors. It is localized most plausibly in MFC, which (with brainstem and striatum) is generally considered as an RL circuit (9,29,30). However, computations in this unit are not used for driving task-related actions, but for driving hierarchically-higher actions, namely to (de)synchronize task modules. This is in line with recent considerations of MFC as a meta-learner (31–34). We tentatively call this unit aMFC, given this region’s prominent anatomical connectivity to autonomous regions (35).

The Control unit was adopted from (16). Its first part contains units that point to specific posterior processing areas, indicating which neurons should be (un)bound. Thus, this area stores the task demands. We labeled this part LFC, given the prominent role of LFC in this regard (36,37). Its second part sends random bursts to posterior processing areas to synchronize currently active areas. Given the prominent anatomical connectivity of pMFC to motor control and several posterior processing areas (35) we tentatively label this part pMFC. The efficiency of this controlling process is largely determined by pMFC theta power: More power leads to more and longer bursts (16). This is consistent with empirical work linking high MFC theta power to efficient cognitive control (26,27). Power in the model pMFC is modulated by the occurrence of negative prediction errors. More specifically, when a negative prediction error occurs, the pMFC node will receive bursts which will increase theta power. In absence of negative prediction errors, this theta power will slowly decrease across trials. This is consistent with the idea that a constant high MFC power might be computationally suboptimal and empirically implausible. For instance, MFC projects to locus coeruleus (LC;(38)); LC firing is thought to be cognitively costly, perhaps because it leads to waste product in the cortex that needs to be disposed (39). In sum, in the Control unit, LFC and pMFC jointly align neural synchronization in modules of the Processing unit to meet current task demands (40,41). Here, the LFC will indicate which modules should be (de)synchronized and the pMFC will exert control over the oscillations in the Processing unit by (de)synchronizing them via random bursts.

Crucially, both Control units use prediction errors, but at a different time scale. The decision of which modules should be synchronized is based on an evaluation of multiple recent prediction errors in the Switch neuron of the RL unit (slow time scale). The pMFC on the other hand will use an evaluation of the last prediction error to evaluate the amount of control that should be exerted (fast time scale). Hence, when an error occurs, the model will initially exert more control on the currently used task module/strategy. If negative prediction errors keep on occurring after the model increased control, it will switch modules/strategies.

Because of its higher plasticity and stability, the full model achieved higher accuracy than the synaptic models. The full model performed better across all learning rates with backpropagation or RBM. With RW, both models showed similar performance for slow learning rates but the synaptic model performed better with fast learning rates. Thus, for simple (linearly separable) learning problems that can be solved very fast, the need for stability is obviated and the advantage of synchronization disappears.

### Experimental predictions

Importantly, our model made several predictions for empirical data. First, it predicts significant changes in the phase coupling between different posterior neo-cortical brain areas after a task switch. Here, we suggest that desynchronization may be important to disengage from the current task. Consistently, (42) found that strong desynchronization marked the period from the moment of disambiguation of ambiguous stimuli to motor responses. Additionally, Parkinson disease patients, often characterized by extreme cognitive rigidity, show abnormally synchronized oscillatory activity (43). Second, we explored midfrontal theta-activation in the time domain by computing the ERP and in time-frequency domain by wavelet convolution. Both analyses showed an increase of theta-power after an error. This was caused by bursts received from the RL unit which elicited a negative peak at approximately 200 ms, corresponding to the FRN (e.g., 19,21,24). Third, we connected the model to research demonstrating theta/gamma interactions where faster gamma frequencies, which implement bottom-up processes, are typically embedded in and modulated by slower theta-oscillations, which implement top-down processes (28,44–46). For this purpose, we considered coupling between pMFC theta phase and gamma amplitude in the Processing unit. Our model predicts a strong PAC increase in the first trial(s) after task switch. This reflects the binding by random bursts control process which is increased after task switches.

### Limitations and extensions

First, because we mainly focused on the biological plausibility and empirical testability of the current model we limited the complexity of the model, especially at its hierarchically higher levels. In its current organization, the model can only determine when a task switch occurred and then make a binary switch to another task module. Hence, the current version of the model can only switch between two task sets. Future work will address this problem by adding second level (contextual) features which allow the LFC to (learn to) infer which of multiple task modules should be synchronized. One useful application of such second level features would be task set clustering, which allows to generalize over multiple contexts. Specifically, if a second-level feature becomes connected to an earlier learned task set, all the task-specific mappings would be immediately generalized to the novel second-level feature. This is consistent with immediate generalization seen in humans (47–49).

Second, our model uses both synchronization and desynchronization which leads to full synaptic gating of task-(ir)relevant modules. It might be suboptimal to always desynchronize all modules that are not currently task-relevant. As suggested by previous work (50), keeping the irrelevant modules at random states (partial gating) might be sufficient to eliminate catastrophic forgetting.

Third, although using negative prediction errors to modulate the control amplitude of the pMFC might be efficient in the current context, this might not be ideal for more complex environments. Thus, a future challenge is combining our model with earlier work that described how a model can (meta-)learn to optimally modulate pMFC activation depending on the environment’s reward and cost structure (32).

Fourth, the model ignored some aspects of oscillatory dynamics. For instance, our model only implements neural synchronization with zero phase lag; yet BBS may be more biologically plausible, and more efficient, with small inter-areal delays (51). Future work will consider an additional (meta-) learning mechanism that learns to synchronize nodes with an optimal phase delay. Additionally, all Processing unit nodes oscillated at the exact same frequency. This scenario might be unrealistic in a typically noisy human brain. Nevertheless, modeling work shows that two oscillators can learn to oscillate at the same frequency via Hebbian Learning (52) in the coupling weights (parameter *C* in equations (1) and (2)). Moreover, this problem is efficiently solved by using a theta-rhythm for delivering the synchronizing bursts, as we implemented here. Specifically, too low-frequency bursts would cause oscillations with (slightly) different (gamma-band) frequencies to drift apart again. With bursts given at a theta-frequency the gamma oscillations have no time to drift apart since the next period of burst occurs before this can happen. In line with this idea, previous work has demonstrated how the model can deal with frequency differences of at least around 2% (16). One might wonder then if the burst frequency could be even higher than theta; however, too high-frequency bursts would result in too noisy signals in the Processing unit. In this sense, theta frequency might strike an optimal balance for guiding gamma oscillations.

### Related work

The current work relies heavily on previous modeling work of cognitive control processes. For instance, in the current model the LFC functions as a holder of task sets which bias lower-level processing pathways (15,53). It does this in cooperation with the MFC. Here, the MFC determines which lower-level task module receives control over behavior (29). The MFC makes this decision based on an RL algorithm (6,9). Hence, the synchronization process in the current model can also be seen as a reinforcement-driven form of synaptic gating (54,55). In biological systems, such gating is plausibly modulated by dopamine. Additionally, also the amount of control/ effort that is exerted in the model is determined by the RL processes in the MFC(31–33). More specifically, negative prediction errors will determine the amount of control that is needed by strongly increasing the MFC signal (29). This is consistent with earlier work proposing a key role of MFC in effort allocation (31,32,56).

In the current model, the MFC thus functions as a hierarchically higher Actor-Critic structure that uses reinforcement learning to estimate its own proficiency in certain tasks. Based on its estimate of the value of a module, and the reward that actually accumulates across trials, it evaluates whether the current task strategy is suited for the current environment. Based on this evaluation, it will decide to stay with the current strategy or switch to another. This is in line with previous modeling work that described the prefrontal cortex as a reinforcement meta-learner (30,33–35).

One problem we addressed in this work was the stability-plasticity dilemma. Previous work on this dilemma can broadly be divided in two classes of solutions. The first class is based on the fact that catastrophic forgetting does not occur when two tasks are intermixed. Thus, one solution is to keep on mixing old and new information (57–60). McClelland et al. (58) suggested that new information is temporarily retained in hippocampus. During sleep (and other offline periods), this information is gradually intermixed with old information stored in cortex. This framework inspired subsequent computational and empirical work on cortical-hippocampal interactions (61–63).

The second class of solutions is based on the protection of old information from being overwritten. Protection can occur at the level of synapses. For example, (64) combined a slow and fast learning system, with slow and fast weights reflecting long- and short-time-scale contingencies, respectively. Another recent idea is to let synapses (meta-)learn their own importance for a certain task (65,66). Weights that are very important for some task are not allowed to change. Hence, information encoded in those weights is preserved. Protection can also be implemented at activation-level. The most straightforward approach to implement such protection is to orthogonalize input patterns for the two tasks (67,68). A broader solution is gating. This means that only a selected number of network nodes can be activated. Because weight change depends on co-activation of relevant neurons (12,69), this approach protects the weights from changing. For example, Masse et al. (50) propose that in each of several contexts, a (randomly selected) 80% of nodes is gated out, thus effectively orthogonalizing different contexts. They showed that synaptic gating allowed a multi-layer network to deal with several computationally demanding tasks without catastrophic forgetting. However, it was unclear how their solution could be biologically implemented. Our solution also exploited the principle of protection. Future work must develop biologically plausible implementations of the mixing principle too and investigate to what extent mixing and protection scale up to larger data sets.

### Summary

We provided a computationally efficient and biologically plausible framework on how neural networks can address the tradeoff between being sufficiently adaptive to novel information, while retaining valuable earlier regularities (stability-plasticity dilemma). We demonstrated how this problem can be solved by adding fast BBS and RL on top of a classic slow synaptic learning network. RL is used to synchronize task-relevant and desynchronize task-irrelevant modules. This allows high plasticity in task-relevant modules while retaining stability in task-irrelevant modules. Furthermore, we connected the model with empirical findings and provided predictions for future empirical work.

## Methods

### The models

As mentioned before and is shown in Figure 1A, our model consists of three units. First, the Processing unit includes the task-related neural network, which is trained with a classical learning rule (backpropagation, Boltzmann, or Rescorla-Wagner). On top of this classical network, an extra hierarchical layer is added where two other units together constitute an Actor-Critic structure (14). The RL unit, adopted from the RVPM (9), functions as Critic and evaluates whether the Processing unit is synchronizing the correct task modules. This evaluation is used by the Control unit (16), which functions as an Actor to drive neural synchronization in the Processing unit. Thus, the Actor-Critic structure allows the models to implement BBS in an unsupervised manner.

#### The Processing unit

An important feature of the current model is that all nodes in the Processing unit consist of triplets of neurons (Figure 5), as in (16). Equations (1)-(5) are taken from (16), but we reproduce them here for readability. Each triplet (node) contains one classical rate code neuron (with activation *x_i_*) which receives, processes and transmits information; and one pair of phase code neurons (*E_i_*, *I_i_*) which organizes processing in the rate code neurons. In line with previous work (16), excitatory neurons are updated by

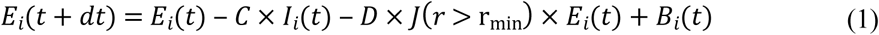

and inhibitory neurons are updated by

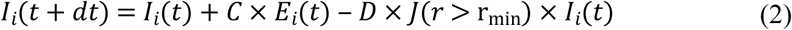

**Figure 5.**
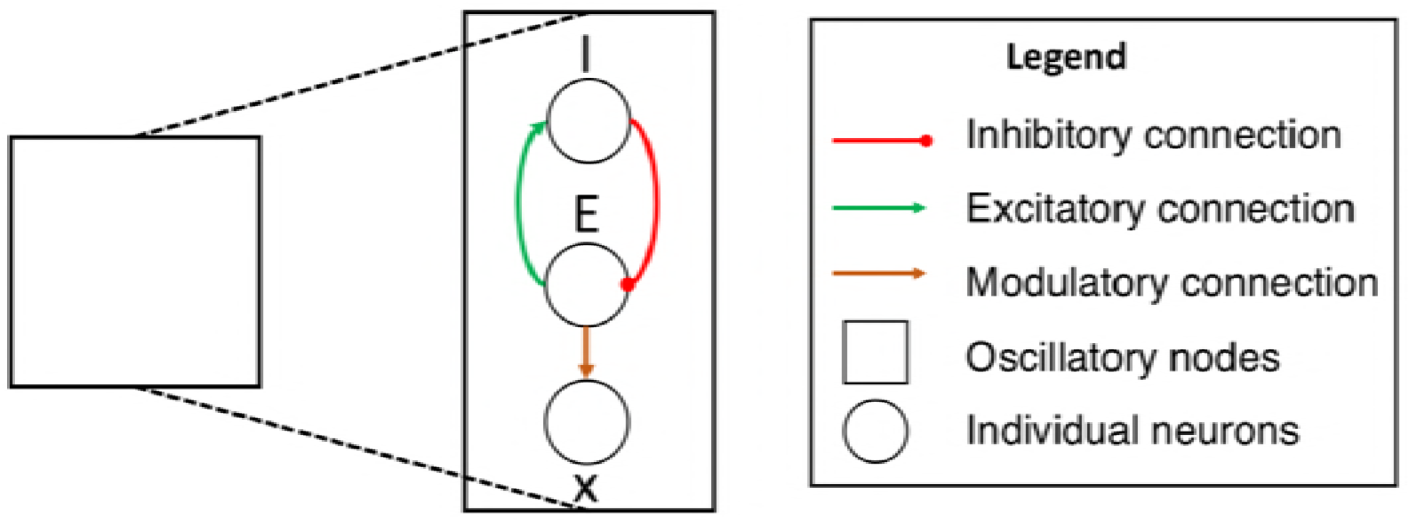
Network node. Illustration of one oscillatory node in the network (see Figure 1C-D), consisting of a triplet of neurons.

The two phase code neurons are thus coupled by a parameter *C*, causing them to oscillate. The strength of the coupling (*C*) determines the frequency of the oscillations, *C*/(2π) (16,70). Task-relevant modules in the processing unit must be bound together. Previous research has proposed that such binding is supported by oscillations in the gamma-frequency band (30-70 Hz; 4). We therefore chose a value for *C* corresponding to a frequency of ~40 Hz. The variable *t* refers to time, and *dt* refers to a time step of 2 msec. The radius (*r* = *E^2^+I^2^*) of the oscillations are attracted towards the value *r*_min_ = 1. This is implemented by the term *D*×*J*(*r>r*_min_)×*E_i_*(*t*) in equation (1) and *D*×*J*(*r>r*_min_)×*I_i_*(*t*) in equation (2). Here, *J*(.) is an indicator function, returning 1 when the radius is higher than the value of *r*_min_ and 0 otherwise. The damping parameter, *D* = .3, determines the strength of attraction. The excitatory neurons of the Processing unit additionally receive a burst,

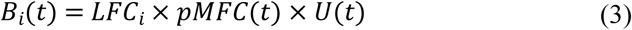

Here, the LFC and pMFC (see Control unit) together determine the burst signal, *B_i_*(*t*), that is received by the excitatory phase code (*E*) neurons. The variable *U*(*t*) is a standardized-Gaussian variable.

The rate code neuron is updated by

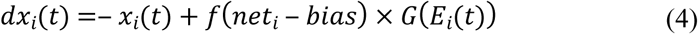

The term -*x_i_*(*t*) will cause fast decay of activation in absence of input. According to this equation, the activation of the rate code neuron at every time step is a function of the net input (*net_i_*) for that neuron multiplied by a function of the excitatory phase code neuron (16),

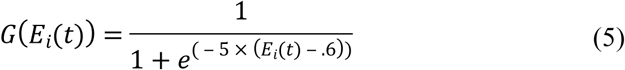

For the multi-layer networks, the rate code neurons have a sigmoid activation function 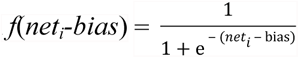. Additionally, these rate code neurons receive a *bias* = 5 to set activation to (approximately) zero in absence of input. In the RW network, the rate code neurons have no bias and follow a linear activation function; *f*(*net_i_ - bias*)*=net_i_*.

Additionally, all weights (*W*) in the Processing unit are subject to learning. Here, learning is done according to one of the three classic learning rules; backpropagation, RBM or RW (10,11,13). A new learning step was executed at the end of every trial. Because activation in the rate code neurons is modulated by *G*(*E_i_*), the activation patterns *x_i_* also oscillate. For simplicity, we use their maximum activation across one trial as input for the learning rules, *X_i_*=max(*x_i_*). Importantly, the standard formulation of the Rescorla-Wagner rule does not combine well with the full model because, in this combination also non-active units would be able learn. To remedy this, a small adjustment was made to the learning rule (13) for the full model. Specifically, we added one term to the classic rule in order to only make co-activated neurons eligible for learning, resulting in

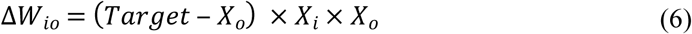

Importantly, this adjustment of the learning rule also results in some costs. First, plasticity decreases because the added term (X_o_) represents the activation of the output unit, which is typically lower than 1 and hence slows down learning. Second, there is a problem at higher learning rates where weights converge to zero and become unable to learn (see dip in Figure 2G). Because the synaptic model obtains no advantage of this adjusted learning rule and we aimed to give the classic model the best chances for competing with the full model, we only used the adjusted learning rule (equation (6)) for the full model.

For the backpropagation and RW networks, a trial ended after 500 time steps (1 sec). Here, the first 250 time steps (500 msec) were simulated as an inter-trial interval in which the Rate code neurons (*x*) did not receive input. In the next 250 time steps, input was presented to the networks. The RBM network also started a trial with 250 time steps without stimulation of the Rate code neurons. After this inter-trial interval the network employs iterations of bidirectional information flow to estimate the necessary synaptic change (11). We used 5 iterations. Every iteration step (2 in one iteration; one step for each direction of information flow) lasted for 250 time steps. The RBM algorithm also employs stochastic binarization of activation levels at each iteration step. Also here, we used the maximum activation over all time steps (*X_i_*) to extract a binary input for that neuron in the next iteration step.

As mentioned in the main text, we compare our new (full) models to models that only use synaptic learning (synaptic models). Thus, those synaptic models only have a Processing unit. Here, all used equations and parameters are the same as described above, except for the synaptic RW model where we use the classic learning rule instead of the one described in equation (6). The only difference is that they do not have phase code neurons and by consequence, *G*(*E_i_*(*t*)) = 1 in equation (5).

*The RL unit.* As RL unit, we implemented the Reward Value Prediction Model (RVPM; Silvetti et al., 2011). Here, there is one expected reward neuron, *V*, which holds an estimation of the reward the model will receive given the task module it used. This estimation is made by

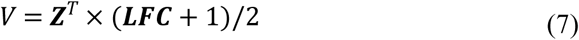

In this equation, ***Z*** is a (column) vector representing the synaptic connections from LFC neurons to the *V* neuron as presented in Figure 1C, D. This vector holds information about the value of specific task modules. Superscript *T* indicates that we transposed the ***Z*** vector. The ***LFC***-term is a vector of LFC values representing which task module drove network behavior on the current trial. These values are normalized, controlling for the fact that LFC neurons can take on negative values. Hence, *V* will represent the expected value of the task module that is synchronized by the LFC represented in the ***Z*** vector. These weights are updated by the RVPM learning rule (9), which is a reinforcement-modulated Hebbian learning rule from the broader class of RL algorithms. All neurons in the RL unit, are rate code neurons which have no time index because they only take one value per trial.

Two prediction error neurons in the RL unit compare the estimated reward (*V*) with the actual received reward. This leads to a negative prediction error *δ*^***-***^ > 0 if the reward is smaller than predicted, *δ*^+^ > 0 if the reward is larger than predicted, and *δ*^***-***^ =*δ*^+^ = 0 if the prediction matches the actual reward (see Silvetti et al. (2011) for more details). The current model accumulates this prediction error signal over several trials to evaluate whether the task rule has changed or not. More specifically, a Switch neuron (*S*) computes a weighted sum of negative prediction errors to determine whether the network is currently using the correct task module. When there is a rapid succession of negative prediction errors, this probably means the task rule has changed. Hence, the network should switch to another strategy. Consequently, activation in the Switch neuron follows

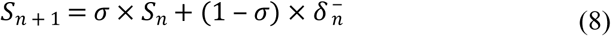

Here, the value of *σ* is set to .8 for the multi-layer models and .5 for the RW model. When activation in this neuron reaches a threshold of .5, it signals the need for a switch to the Control unit (see also equation (12)) and resets its own activation to zero. In the equation, *n* refers to the trial number.

#### The Control unit

*A*s in previous work (16), the Control unit consists of two parts, corresponding to posterior medial (pMFC) and lateral (LFC) parts of the primate prefrontal cortex.

The modelled pMFC represents one node (Figure 5) consisting of one phase code pair (*E*_pMFC_, *I*_pMFC_) and a rate code neuron (*pMFC*). The phase code neurons obey the same updating rules as given by equation (1) and (2). In the pMFC, which executes top-down control, the value of *C* is such that oscillations are at a 5Hz (theta-) frequency, in line with suggestions of previous empirical work (26,27). Since a constant high MFC power is computationally suboptimal and empirically implausible (39), the radius of the pMFC was attracted towards a small radius, r*_min_*=.05. The damping parameter was set to *D* = .03, in order to let the amplitude of the pMFC oscillations decay slowly over trials. The burst signal of the pMFC was determined by the negative prediction error signal of the previous trial,

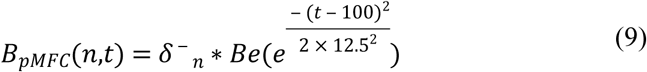

Here, the burst signal at one time point in one trial is determined by the size of the negative prediction error at the previous trial and a Bernoulli process Be(p(t)) which is one with probability P(t). The probability P(t) corresponds to a Gaussian distribution over time that has its peak at 100 time steps and a standard deviation of 12.5 time steps, representing a delay of communication between the pMFC and the RL unit. Hence, when the previous trial elicited a negative prediction error, bursts are sent to the excitatory neuron of the pMFC. Consequently, these bursts have the size of the negative prediction error and are most likely to occur at 100 time steps (200 ms) after feedback. This burst signal will increase the amplitude of the pMFC phase code neurons when a negative prediction occurs, after which it will again slowly decay towards r_min_.

In line with the previous study (16), activation in the rate code neuron of the pMFC follows

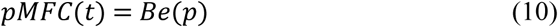

Again, this equation represents a Bernoulli process *Be*(*p*) which is 1 with probability *p*. The probability

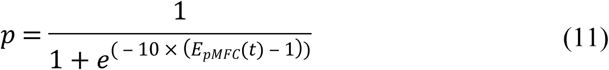

is a sigmoid function which has its greatest value when the *E_pMFC_*_(*t*)_ is near its top and its amplitude is sufficiently strong. Hence, every time the oscillation of the *E_pMFC_*-neuron reaches its top, the probability of a burst becomes high. Thus, bursts are phase-locked to the theta oscillation, implying that the pMFC determines the ‘when’ of the bursts (see (16) for more details).

In general, the model implements a “win stay, lose shift” strategy, shifting attention in LFC when reward appears less than expected. As shown in Figure 1C, D, the LFC consists of three rate code neurons that each have a pointer to one (or two) of the different modules in the Processing unit. One of these LFC neurons is connected to the visible layers (input and output) for the multi-layer networks and the input layer for the RW network and has a constant value of 1. Each of the other two LFC neurons are connected to one of the two modules in the hidden, or in the case of the RW network, the output layer. For these neurons, at trial *n* = 1 a random choice is made where one neuron is set to 1 and the other to −1. In trials *n* > 1, they obey

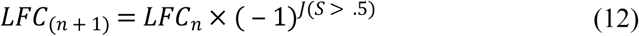

Hence, the network always synchronizes one task module with the in- and output layers and desynchronizes the other task module. When the Switch neuron, *S,* reaches the threshold, indicator function *J*(.) will return 1 instead of 0. This will change the sign in both LFC neurons connected to the task modules and therefore synchronize the previously desynchronized module and vice versa. This can easily be scaled up to more modules and more task rules by letting the model make a random choice or including context-specific input.

### The task

We test our model on a reversal learning task (71,72). We divide the task in three equally long parts. In the first two parts, the model should learn two different new task rules (rule 1 and rule 2 in parts 1 and 2, respectively). In the third part, the model has to switch back to following rule 1.

In the context of the multi-layer networks, we chose a Stroop-like task consisting of 2400 trials in total. Stimuli contain three crucial features. They are words (“red” or “blue”) printed in a certain color (red or blue) and style (bold or italic). There are two response options. The task is to respond to the word when it is printed in bold and to the color when it is printed in italic. During rule 1 they should respond with R1 for red and R2 for blue. This is reversed for rule 2. All stimuli are presented equally often in random order.

For the RW network, which cannot handle such complex task rules, we use simple S-R associations as task rules. According to rule 1, R1 leads to reward after presentation of S1 and R2 leads to reward after presentation of S2. For rule 2 these associations are reversed, linking R1 with S2 and R2 with S1. Here, the task is divided in three parts of 80 trials each, making a total of 240 trials. Again, in each part, each possible stimulus is presented equally often in random order.

### Simulations

To test the generality of our findings, we varied the synaptic learning rate. This parameter was varied from 0 to 1 in 11 steps of .1. For each value, we performed 10 replications of the simulation. In every simulation, the strength of synaptic connections at trial 1 was a random number drawn from the uniform distribution, multiplied by the bias value (and 1 for the RW based model).

The effects of other model parameters were already demonstrated in previous work (9,16), but we again validated that the model shows qualitatively similar patterns when we varied some of the parameters. This was true when we changed the frequency *C*/2π of oscillations in the Processing unit to 30 Hz; attracted the pMFC amplitude to a value r_min_=.5; used a Switch threshold of .45 or .55 in equation (12); or varied the learning rate in the RL unit.

### Statistical analyses

For the purpose of comparison, we divided the trials of the task for every model into 60 bins. For the RW based model, bin size equals 4 trials; for the multi-layer models, bin size equals 40 trials. We evaluate the performance of our model on several levels. First, we evaluate overall task accuracy. Second, we evaluate plasticity. For this purpose, we explore the performance of the model right after the switch from task rule 1 to task rule 2; we compute the mean accuracy on the first 5 bins after the switch. Third, we evaluate stability. In particular, we explore the interference of learning task rule 2 in between two periods of performing on task rule 1. For this purpose, we compare the accuracy right after (5 bins) the second switch and right before the first switch (5 bins). If the model saved what it has learned about task rule 1, this difference should be zero. If the model displays catastrophic forgetting it would have a negative stability score.

Importantly, we also connect with empirical data and describe testable hypotheses for future empirical work. Because the multiple iterations performed by the RBM algorithm render it more complex to extract the oscillatory data, and because this algorithm is less biologically plausible, we focused these analyses on the backpropagation and RW model. As a measure of phase synchronization between excitatory neurons in the Processing unit, we compute the correlation at phase lag zero. A correlation of 1 indicates complete synchronization and −1 indicates complete desynchronization. Phase-amplitude coupling (PAC) is computed as the debiased phase-amplitude coupling measure (dPAC; 25) in each trial. Here,

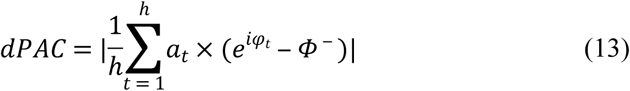

in which

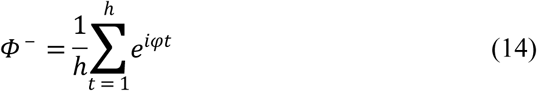

In these equations, *t* represents one time step in a trial, *h* is the number of time steps in a trial, *a* is the amplitude, *φ* is the phase of a signal, and *i*^2^ = −1. In the current paper, we are interested in the coupling between the phase of the theta oscillation in the pMFC node of the Control unit and the gamma amplitude in the Processing unit. Phase was extracted by taking the analytical phase after a Hilbert transform. The gamma amplitude was derived as the mean of the excitatory phase code activation of all nodes in the Processing unit by

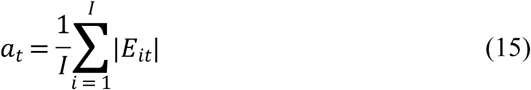

with *I* being the number of nodes in the Processing unit, *t* referring to time and *E_i_* being the respective excitatory phase code neuron.

For all measures, we represent the mean value over *Nrep* = 10 replications and error bars or shades show the confidence interval computed by mean± 2*×*(SD/√Nrep).

Additionally, we evaluated the pMFC theta activation. First, in order to illustrate the bursts described in equation (10), we computed the ERP during the intertrial interval after error trials. Second, we evaluated power in the time frequency domain. Time–frequency signal decomposition was performed by convolving the signal (e.g., for an *E* neuron) by complex Morlet wavelets, 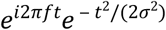, where i^2^ =-1, t is time, *f* is frequency, ranging from 1 to 10 in 10 linearly spaced steps, and σ=4/(27πf) is the “width” of the wavelet. Power at time step t was then computed as the squared magnitude of the complex signal at time t and frequency f. We averaged this power over all simulations and all replications of our simulations. This power was evaluated by taking the contrast between the inter-trial intervals following correct (1) and error (0) reward feedback.

### Data and software availability

Matlab codes that were used for both the model simulations and data analysis are available on GitHub (https://github.com/CogComNeuroSci/PieterV_public). We will also provide adapted versions of this code for use with Python.

## Acknowledgements

The current work was supported by grant BOF17/GOA/004 from the Ghent University Research Council. PV was also supported by grant 1102519N from Research Foundation Flanders. We thank Daniele Marinazzo and Cristian Buc Calderon for helpful comments.

